# A cell-free DNA metagenomic sequencing assay that integrates the damage response to infection

**DOI:** 10.1101/648592

**Authors:** Alexandre Pellan Cheng, Philip Burnham, John Richard Lee, Matthew Pellan Cheng, Manikkam Suthanthiran, Darshana Dadhania, Iwijn De Vlaminck

## Abstract

High-throughput metagenomic sequencing offers an unbiased approach to identify pathogens in clinical samples. Conventional metagenomic sequencing however does not integrate information about the host, which is often critical to distinguish infection from infectious disease, and to assess the severity of disease. Here, we explore the utility of high-throughput sequencing of cell-free DNA after bisulfite conversion to map the tissue and cell types of origin of host-derived cell-free DNA, and to profile the bacterial and viral metagenome. We applied this assay to 51 urinary cfDNA isolates collected from a cohort of kidney transplant recipients with and without bacterial and viral infection of the urinary tract. We find that the cell and tissue types of origin of urinary cell-free DNA can be derived from its genome-wide profile of methylation marks, and strongly depend on infection status. We find evidence of kidney and bladder tissue damage due to viral and bacterial infection, respectively, and of the recruitment of neutrophils to the urinary tract during infection. Through direct comparison to conventional metagenomic sequencing as well as clinical tests of infection, we find this assay accurately captures the bacterial and viral composition of the sample. The assay presented here is straightforward to implement, offers a systems view into bacterial and viral infections of the urinary tract, and can find future use as a tool for the differential diagnosis of infections.

## INTRODUCTION

Differential diagnosis of infectious disease in humans is complex. Metagenomic high-throughput DNA sequencing offers an unbiased approach for the detection of pathogens in clinical samples^1–4^, but the presence of a pathogen is not necessarily synonymous with disease^5^. Some microbes are commensals in all human hosts, some only cause disease in some hosts, and others cause disease in all hosts. To bring clarity to the lexicon of microbial pathogenesis, Casadevall and Pirofski defined infectious disease as a clinical manifestation of damage to the host that results from host-microbe interaction^5,6^. In this framework, the degree of host damage, mediated by the host response and/or by the pathogen, offers a quantifiable metric that can be used to distinguish between different outcomes of infection^6^.

We report a high-throughput metagenomic sequencing assay that can both detect a diverse array of bacterial and viral pathogens and quantify damage to host tissues. The assay implements whole-genome bisulfite sequencing (WGBS) of cell-free DNA (cfDNA), small fragments of DNA released by host or microbial cells into blood, urine and other bodily fluids, and brings together two previously reported concepts. First, the assay implements a genome-wide measurement of cytosine methylation marks comprised within cfDNA – marks that are highly cell, tissue and organ-type specific – to determine the cell and tissue types that contribute to the mixture of host cfDNA in a sample. Several recent studies have shown that profiling CpG methylation marks in urinary or plasma cfDNA, via whole-genome sequencing, targeted sequencing, or PCR assays, can be used to determine their tissues-of-origin and to quantify tissue-specific injury in various diseased settings ^7–9^. Here, we explore this concept for the monitoring of injury due to infection. Second, the assay quantifies the relative abundance of microbes via WGBS of cfDNA. Several studies have investigated the utility of conventional, metagenomic sequencing of cfDNA for infection testing in clinical samples^2,4,10,11^. We show here that WGBS is compatible with such analyses.

We investigated the utility of this assay to monitor infectious complications of the urinary tract after kidney transplantation. More than 80,000 patients receive lifesaving kidney transplants worldwide each year^12^. Immunosuppression after transplantation is required to manage the risk of rejection but leaves patients vulnerable to viral and bacterial infection. BK Polyomavirus (BKV) infection has emerged as serious risk factor for allograft survival. BKV reactivation occurs in up to 73% of kidney transplant recipients, and leads to BK Polyomavirus Nephropathy (BKVN) in up to 8% of patients^13,14^. Renal biopsies are currently required to confirm BKVN and to distinguish BKVN from BKV reactivation without nephropathy (BKV+/N-). While BKVN histology is characterized by inflammation and necrosis of tissue, biopsies from BKV+/N-patients are similar to those without reactivation^13^. It remains unclear whether BKV reactivation alone induces kidney damage. Bacterial urinary tract infection (UTI) affects approximately 43% of kidney transplant recipients in the first 42 months post-transplant^15^. There is a disagreement in the literature regarding the appropriate balance between mitigating the risks of infectious complications and adverse effects of antimicrobial treatment for UTI. In this study, we describe a urinary cfDNA assay that can identify viral and bacterial infectious agents and can quantify the degree of host injury related to UTI.

## RESULTS

### Methylation marks are cell, tissue and organ type specific

We performed WGBS (**Fig. 1a**) on 51 urinary cfDNA isolates collected from a cohort of kidney transplant recipients (**Fig.1b**) and used computational methods to quantify the burden of viral and bacterial cfDNA and the cell and tissue types of origin of host-derived cfDNA (**Fig.1c**). We assayed urinary cfDNA isolates from patients who had a same-day corresponding bacterial culture (UTI positive, UTI group, n=12; UTI negative, no-UTI group, n=12), and from patients that were BKV positive in the blood and confirmed to have BKVN by biopsy (BKVN, n=9), BKV positive in the blood without evidence of BKVN on biopsy (BKVN+/N-, n=7) and negative for BKV in the blood and with normal surveillance biopsy (Normal, n=6). In addition, we analyzed urinary cfDNA obtained from patients within the first three days after transplantation (Early group, n=5). To obtain sequence information after bisulfite conversion of these molecules, we used a single-stranded sequencing library preparation^1,16^ (**Fig. 1a**). This library preparation employs ssDNA adapters and bead ligation to create diverse sequencing libraries from short, highly fragmented cfDNA^16,17^. We obtained 104.5 +/- 43 million paired-end reads per sample, corresponding to a per-base human genome coverage of 1.4-4.1x (see Methods).

**Figure 1.**
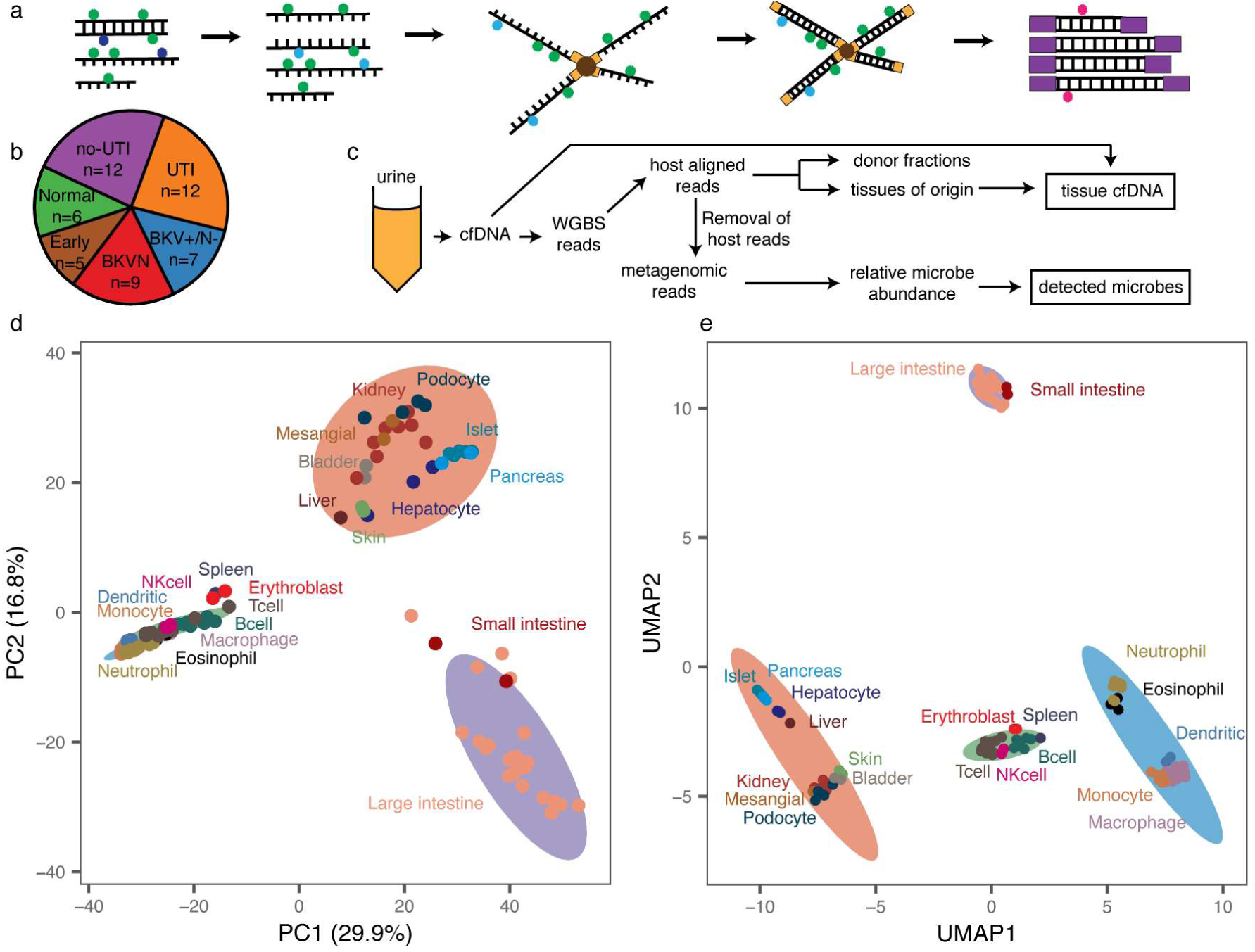
Study methodology and cell type specificity of genome-wide methylation profiles. **a** Schematic of single-stranded library preparation method. cfDNA is denatured and treated with sodium bisulfite, which converts unmethylated cytosines (dark blue) into uracils (light blue) but not methylated cytosines (green). Bisulfite-treated DNA is first ligated to single-stranded adapters and bound to magnetic beads. Second-strand synthesis, and double-stranded adapter ligation are performed on the beads. The final step is a PCR, which converts uracils to thymines (red). **b** Pie chart with summary of samples included in this study, colored by pathology. **c** Schematic of WGBS analysis workflow. **d** Principal component analysis of reference whole-genome methylation profiles from human tissues. **e** Uniform manifold approximation projection of reference methylation profiles. Ellipses in **d, e** are normal 95% confidence ellipses (K-means, 4 centers).

We sought to implement a reference-based approach for cell-type deconvolution, thereby taking advantage of the large and growing number of genome-wide methylation profiles of tissues and cell types of interest that are available in public repositories. We downloaded 112 genome-wide methylation datasets representing 16 different tissue types (**Supplemental table 1**)^18–22^, and determined tissue-specific differentially methylated regions (DMRs) using Metilene^23^. We compared CpG methylation profiles of each tissue group in a one-versus-one approach and found 91,275 DMRs, with an average length of 453 base pairs (bp). Principal component analysis (PCA) of the methylation density measured across all DMRs revealed global tissue-specific clustering, with three heterogeneous clusters representing blood, gut and a diverse group of other solid organ tissues (**Fig.1d, Supplemental Fig. 1**). To identify both global and local structural features within reference methylomes, we applied uniform manifold approximation projection (UMAP)^24^. We found that UMAP further resolves reference methylation profiles into clusters comprised of specific cell-types (**Fig. 1d,e**). For example, among myeloid cells, cell types with similar lineages such as macrophages and monocytes clustered more closely than those from other lineages on the UMAP projection. These analyses confirm that genome-wide methylation profiles are cell, tissue and organ-type specific, and can in principle inform its tissue of origin, as described previously^7,8,25–27^.

### Cell-free DNA origin associated with infection

To determine methylation marks comprised within urinary cfDNA, we first aligned the WGBS reads to a human reference genome via bwa-meth^28^. We quantified the efficiency of bisulfite conversion achieved in experiments from the fraction of reported methylated C[A/T/C] base pairs, which are rarely methylated in humans^29^. We found a conversion efficiency greater or equal to 94.5% for all cfDNA isolates (see Methods). We next projected the genome-wide cfDNA CpG methylation profiles onto the two-dimensional feature spaces generated by PCA and UMAP for the 112 public references. We found that cfDNA profiles organized between the cluster that comprised kidney tissue and the white blood cell cluster on the PCA and UMAP two-dimensional projections (**Fig. 2 a,b**). This observation provided a first indication that urinary cfDNA originates primarily from blood cell types and kidney tissue. Other cell types found in the urinary tract contributed less significantly to urinary cfDNA.

**Figure 2.**
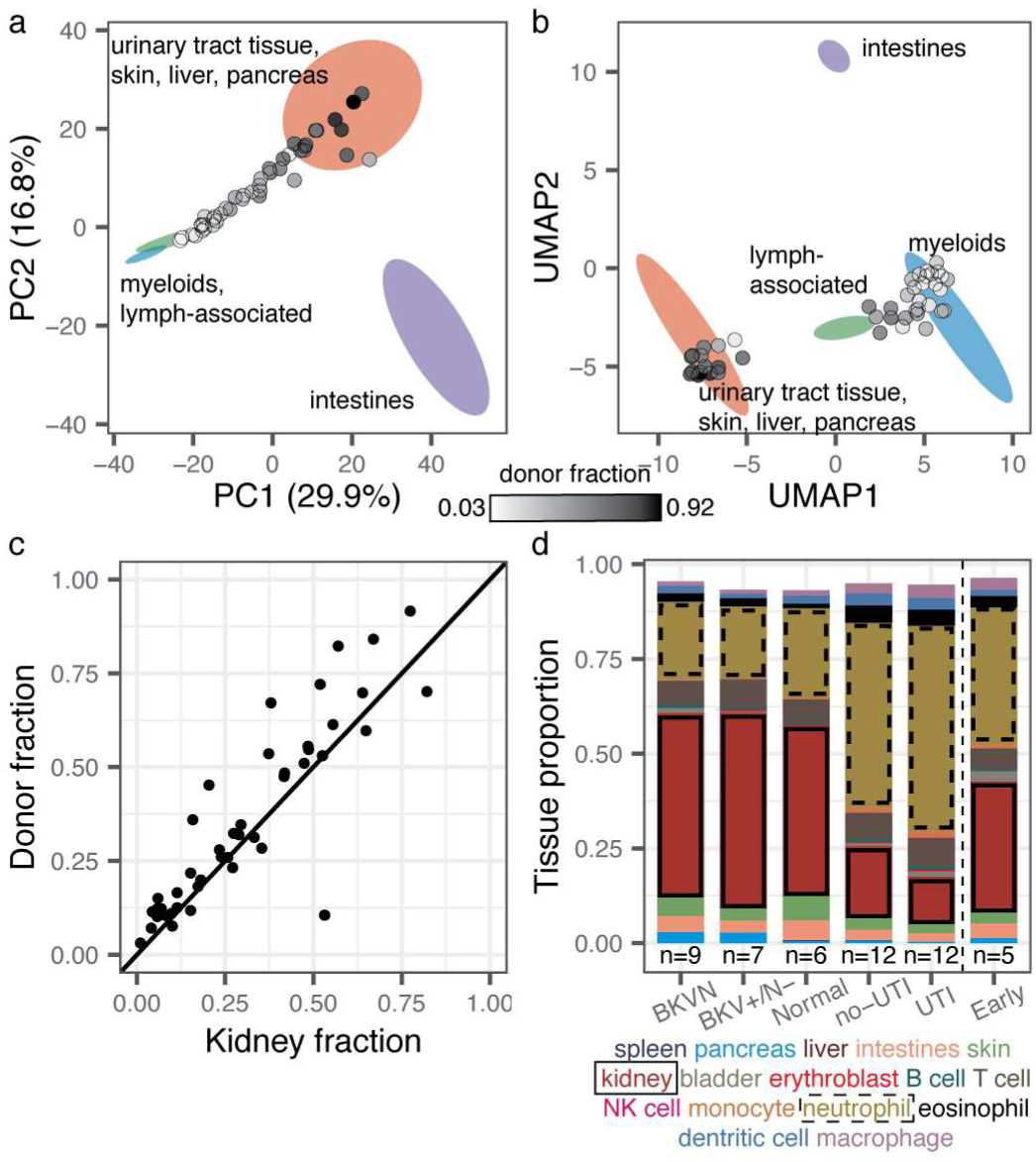
Methylation marks within cfDNA inform its tissues-of-origin. **a, b** Projection of cfDNA genome-wide methylation profiles onto PCA and UMAP feature space (from Fig. 1d,e). Data points colored according to donor fraction (n=44). **c** Comparison of the proportion of kidney-derived cfDNA and donor-derived cfDNA measured for sex-mismatched donor-recipient pairs (n=44). Spearman’s rho=0.88, p-value < 2.2×10^-16^. Samples from dual bone-marrow and kidney transplants are excluded. **d** Barplot of cfDNA cell and tissue type composition measured for clinical groups.

We next computed the proportion of transplant donor-derived cell-free DNA (ddcfDNA) in these samples. We and others have identified ddcfDNA as a non-invasive, quantitative marker of graft injury in solid-organ transplantation^2,4,30,31^. For sex-mismatched donor-recipient pairs, the proportion of ddcfDNA can be estimated by evaluating the coverage of the Y chromosome relative to the autosomal chromosomes (see Methods)^2,32^. We verified that the proportion of ddcfDNA measured by sequencing of bisulfite-treated cfDNA matched the proportion of ddcfDNA measured using conventional sequencing (n=36 matched samples, Spearman’s rho = 0.97, p-value < 2.2×10^-16^, see Methods, **supplemental figure 2**), and then quantified the proportion of ddcfDNA in urine for all sex-mismatched donor-recipient transplant pairs (n=46). We observed a very large range of ddcfDNA values across all samples (3%-99%). Superimposing the ddcfDNA proportion on the PCA and UMAP projections revealed that samples with a higher proportion of ddcfDNA landed closer to the reference cluster comprised of kidney tissue (**Fig. 2a,b**, n=44, two samples from a patient that received both a kidney and bone marrow transplant were excluded from this analysis, because the donor fraction represents the summation of kidney DNA and engrafted bone marrow-derived cells for this case). This observation provided a second line of evidence that urinary cfDNA is a mixture of cfDNA derived from blood cell types and kidney tissue.

To quantify the contributions of different tissues to the mixture of cfDNA in urine, we implemented quadratic programming. Quadratic programming retrieves the fractional contribution of each tissue, π_i_, from the ensemble cfDNA methylation profile Y, and the public reference methylation profile for each tissue, X_i_: Y=π_i_X_i_+ε, where ε is an error term (see Methods)^33^. Using this approach, we found an excellent quantitative agreement (Spearman’s rho=0.88, p-value < 2.2×10^-16^, **Fig. 2c**) between the proportion of kidney specific cfDNA (determined from methylation marks) and ddcfDNA (determined from genetic marks in cfDNA) for sex-mismatched donor-recipient transplant pairs (n=44). This analysis provided support for the use of our bioinformatic and molecular approaches to quantify the tissue and cell types of origin of cfDNA in urine.

We proceeded to analyze the relative contributions of all cell and tissue types comprised within the pure cell and tissue references against clinical tests of infection. We found that the cfDNA cell and tissue composition was associated with infection status (**Fig. 2d**). For example, the relative contribution of kidney-derived cfDNA was elevated in samples from patients with BKV infection compared to patients diagnosed with bacterial UTI (p-value = 2.0×10^-4^, mean of 48.6%; mean 12.5%, respectively, **Fig. 2d**). We further found that leukocytes were enriched in samples from patients diagnosed with bacterial UTI by conventional culture, and that neutrophils are the main contributors to the differences in white blood cell content, as expected from their role as first responders to infection (**Fig. 2d**)^34^.

The relative proportion of cfDNA from a specific tissue is a function of the proportion of DNA released from all other cell types or tissues and may therefore be a convoluted measure of tissue-specific injury. To overcome this limitation, we computed the absolute concentration of tissue-specific cfDNA by multiplying the proportion of tissue and cell type specific DNA obtained using the approaches above with the concentration of total host-derived cfDNA in the sample (see Methods). We observed marked temporal dynamics of the concentration of cfDNA from different cell and tissue types in absence of infection (no-UTI, Normal and Early groups, **Fig. 3a**). Recovery of postoperative stress in the first three days after transplantation was associated with a marked increase in cfDNA from most tissues. This signal of early post-operative injury decayed to a low baseline within 10 days after transplantation, and after three months post transplantation quiescence was observed with markedly low amount of cfDNA from all cell and tissue types in absence of infection.

**Figure 3.**
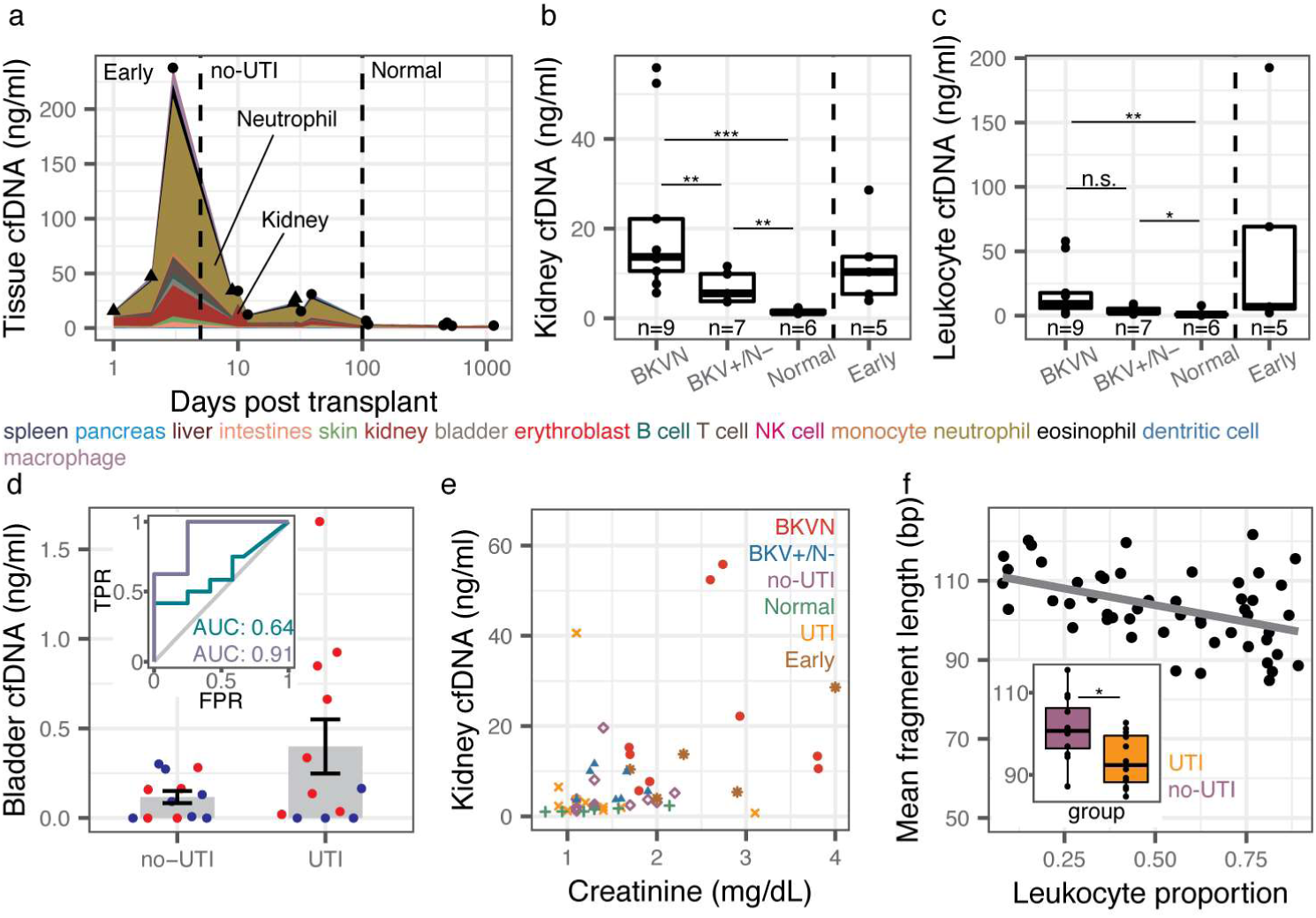
Cell-type deconvolution of urinary cfDNA reveals host response to infection. **a** Concentration of tissue-specific cfDNA for samples with no clinical indication of infection (triangles indicate mean for multiple measurements at the same time point, single samples indicated with circles). **b,c** Kidney (b) and leukocyte (c) cfDNA concentration for BKVN, BKV+/N-, Normal and Early groups. **d** Bladder cfDNA concentration for no-UTI and UTI groups. Inset shows receiver operating characteristic analysis of the performance of bladder cfDNA in distinguishing no-UTI from UTI groups (blue) and in distinguishing hematuria (RBC per HPF >4) from no hematuria (RBC per RBC ≤4) in samples from the UTI group. **e** Serum creatinine versus kidney urinary cfDNA concentration (n=50). Spearman’s rho=0.51 p-value=1.4×10^-4^. **f** Mean host cfDNA fragment length versus leukocyte proportion (n=51). Spearman’s rho=-0.45, p-value=1.1×10^-3^. Inset shows boxplots of fragment lengths measured for the UTI and no-UTI groups. Boxplot features detailed in the Methods section. * p-value< 0.05, ** p-value< 0.01, *** p-value< 0.001.

We examined whether the cfDNA concentration of certain tissues was associated with infection pathology. We first examined all samples from patients that were screened for BKVN via needle biopsy (all samples collected after day 100). We observed marked differences in the concentration of kidney-derived cfDNA for samples from patients diagnosed with BKVN (BKVN vs Normal, mean kidney cfDNA 21.9 ng/ml, and 1.5ng/ml, respectively p-value = 4.0×10^-4^, **Fig. 3b**), indicating significant tissue injury associated with BKVN. In addition, we found that this cfDNA measurement can distinguish BKVN from BKV reactivation without nephropathy (BKVN vs BKV+/N-, mean kidney cfDNA 21.9 ng/ml and 6.9 ng/ml, respectively, p-value = 7.9×10^-3^), and BKV+/N-from Normal (mean kidney cfDNA 6.9 ng/ml and 1.5ng/ml, respectively, p-value = 1.2×10^-3^). The concentration of leukocyte cfDNA was elevated in urine from patients diagnosed with BKVN (mean 19.0 ng/ml vs. 1.9 ng/ml in Normal, p-value = 2.8×10^-3^) and BKV+/N-(mean 4.0 ng/ml vs. 1.9 ng/ml, p-value = 3.5×10^-2^) but could not distinguish BKVN from BKV reactivation without nephropathy (p-value = 5.5×0^-2^) (**Fig. 3c**). Together these experiments point to the utility of the assay presented here to non-invasively distinguish nephropathy and inflammation due to BK virus infection. The kidney cfDNA concentration in samples collected from patients within the first three days after transplantation, a period during which we observed significant post-operative injury (**Fig. 3a**), was also elevated (12.4 ng/ml) and could not be differentiated from the concentration measured in samples from patients with BK disease (p-value=0.30 and 0.20 when compared to BKVN and BKV+/N-, respectively). This last observation underlines the importance of paired metagenomic evaluation to distinguish infection and non-infection related host injury.

We next evaluated all samples from patients that were screened for UTI via conventional urine culture (no-UTI group and UTI group, n=12 each). These samples were collected between days 8 and 55 post-transplant, a period in which we observed significant host injury in absence of infection (**Fig. 3a**), and were evaluated for hematuria by microscopy, a clinical marker of injury. We found that both bladder and leukocyte-derived cfDNA were elevated in samples from patients diagnosed with bacterial UTI and hematuria (Red Blood Cell [RBC] counts per High Power Field [HPF] greater than 4) compared to patients diagnosed with UTI in the absence of hematuria (**Fig. 3d**, receiver operator characteristic analysis, area under the curve [AUC] = 0.91 for bladder cfDNA). We further found correlations between red blood cell (RBC) counts and the concentrations of bladder cfDNA in urine (all samples for which RBC per HPF was measured, n=24, Spearman’s rho = 0.43, p-value = 3.5×10^-2^). Together, these observations demonstrate the utility of our assay to assess the severity of injury due to bacterial UTI. The performance of the concentration of bladder cfDNA in distinguishing bacterial UTI and absence of UTI with and without hematuria was modest (AUC = 0.64 for bladder cfDNA, **Fig. 3d**), which is likely explained by the significant non-infection related injury in this patient population in the sample time window.

Last, we asked whether the concentration of kidney derived cfDNA correlated with serum creatinine, a clinical marker of kidney function. Creatinine is a waste product of muscle metabolism and elevated serum creatinine levels are an indication of poor kidney function. We found good agreement between kidney-specific cfDNA in urine and serum creatinine (all samples for which serum creatinine was measured, n = 50, Spearman’s rho=0.51, p-value=1.5×10^-4^). Together, the data presented in Figure 3 provide strong support for the use of WGBS of cfDNA to quantify host injury due to viral and bacterial UTI.

The pattern of degradation of cfDNA in plasma has previously been shown to depend on their origin and pathology. For example, tumor-derived cfDNA was found to be significantly shorter than cfDNA from normal tissue^35,36^. Here, we sized urinary cfDNA using paired-end read mapping (see Methods). Bisulfite treatment is known to lead to DNA degradation^37^, and this was corroborated by our cfDNA sizing assay. We found that bisulfite-treated cfDNA is on average 10.1 bp shorter than untreated cfDNA (n = 38 matched samples). We furthermore found that the mean fragment length of host-derived cfDNA was negatively correlated with the white blood cell proportion (Spearman’s rho = −0.45, p-value = 1.1×10^−3^, n=51) and was shorter for samples from patients with bacterial infection than from patients without bacterial UTI (p-value = 2.4×10^-2^, mean length of 93 and 101bp, respectively, **Fig. 3f, Supplemental Fig. 3**). This observation is in line with previous reports that leukocytes create a degradative environment for DNA^38,39^. The fragment size profile of cfDNA may offer an additional metric by which patients with different infectious pathologies can be stratified.

### Whole genome bisulfite sequencing of cfDNA identifies clinically-relevant pathogens

Bisulfite conversion of DNA followed by PCR converts unmethylated cytosines into thymines. A corollary of bisulfite treatment is a decrease in cytosine content, and a reduction in overall read complexity. To determine whether WGBS can be used to identify specific uropathogens despite the reduction in sequence complexity inherent to bisulfite conversion, we compared pathogen abundances measured after shotgun sequencing of bisulfite-treated and untreated cfDNA (matched samples, n=38). We determined the relative representation of bacteria and viruses in these datasets, using approaches previously described^2,3,40^ (see Methods). We computed the representation of microbial genomes relative to human genome copies and expressed this quantity as relative genome equivalents (RGE, see Methods). Figure 4a shows a close quantitative agreement between the species abundance measured for bisulfite-treated and untreated cfDNA, confirming that it is possible to broadly identify microbial cfDNA via shotgun sequencing of bisulfite-treated cfDNA (**Fig. 4a**, Spearman’s rho = 0.72, p-value < 2.2×10^-16^).

**Figure 4.**
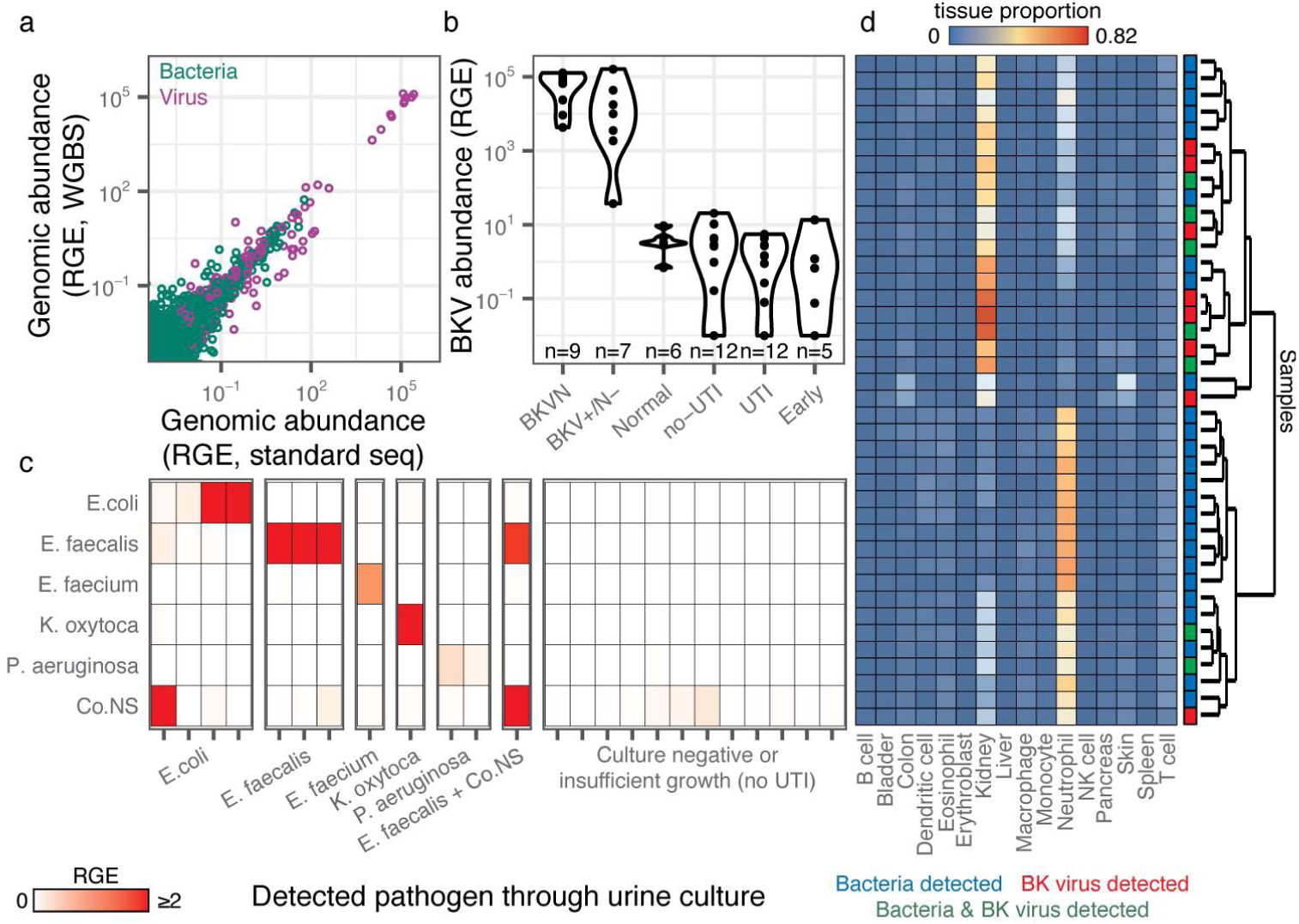
WGBS of microbial cfDNA for clinical pathogen identification. **a** Scatterplot of relative genomic abundance of bacteria (green) and viruses (purple) measured by WGBS and conventional cfDNA sequencing. Spearman’s rho = 0.72, p-value < 2.2×10^-16^. Each data point represents the genomic abundance of a single microbe in matched bisulfite and untreated urinary cfDNA. **b** Violin plots of BKV sequence abundance in all samples. c Relative genomic abundance of microbes identified through urine culture. **d** Heatmap of tissue proportions in samples where a microbe was detected through WGBS. Rows are hierarchically clustered according to tissue composition.

We assessed the relative genomic abundance of cfDNA from bacterial and viral pathogens identified by conventional diagnostic assays. In 9 out of 9 BKVN samples and 6 out of 7 BKV+/N-samples, we identified high BK viral loads (RGE > 10^3^, **Fig. 4b**). These values correlated strongly with matched plasma BKV copies as determined by quantitative PCR (Spearman’s rho=0.81, p-value=6.2×10^-6^). We next compared the relative genomic abundance of bacterial cfDNA for patients diagnosed with bacterial infection (12 samples matched with a positive clean-catch midstream urine culture, UTI group), to the relative genomic abundance measured for 12 negative clean-catch midstream urine cultures (no-UTI group). Of these positive urine cultures, 11 had a single identifiable bacterium (*Escherichia coli*, n=4; *Enterococcus faecalis*, n=3; *Pseudomonas aeruginosa*, n=2; *Enterococcus faecium*, n=1; *Klebsiella oxytoca*, n=1), while a single sample presented 2 different bacterial species (*E. faecalis* and coagulase-negative Staphylococcus). We found agreement between the urinary cfDNA assay described here and conventional bacterial culture (detection accuracy = 100%, no information rate = 16.0%, p-value [accuracy > no information rate] < 2.2×10^-16^, **Fig. 4c**). These results support the use of WGBS of cfDNA as an assay to screen for potential pathogens in clinical isolates.

Finally, we tested whether the cfDNA cell and tissue-type composition depended on the presence or absence of viral and bacterial pathogens as determined by WGBS. We used unsupervised hierarchical clustering of the cfDNA cell and tissue type composition for all samples in which BK virus (RGE > 10^3^) or a potential bacterial uropathogen was detected (RGE > 0.09, the lowest corresponding relative genomic abundance observed in the comparison to clinical metrics described above). This analysis, shown in figure 4d, summarizes the major layers of information that are made accessible with the cfDNA assay reported here, and shows that the cfDNA tissue and cell type composition is associated with the presence or absence of viral or bacterial uropathogens.

## DISCUSSION

We have described a metagenomic assay that simultaneously quantifies the abundance of a large array of viruses and bacteria in clinical samples, and the degree of host injury. This work is motivated by the need to integrate information about host-microbe interactions in clinical metagenomic assays in order to distinguish infection from infectious disease, and to assess the severity of disease. The assay reported here takes advantage of genome-wide profiling of CpG methylation marks comprised within cfDNA to quantify the contributions of different cell and tissue types to the mixture of cfDNA in the sample, and thereby the degree of host damage. Compared to conventional metagenomic cfDNA assays^2,41^, this assay requires a single additional experimental step, bisulfite treatment of the cfDNA isolate, which is inexpensive and can be completed within approximately two hours.

We tested the utility of this assay to monitor viral and bacterial infections of the urinary tract in a cohort of kidney transplant recipients. We found that the concentration of cfDNA derived from different cell and tissue types was a function of infection status in these patients. Patients diagnosed with BKVN had elevated kidney-specific cfDNA in urine compared to a Normal control group. BKV reactivation without nephropathy was also characterized by elevated levels of kidney-specific DNA but not to the same degree as BKVN. Our findings suggest that there may be kidney damage occurring before the onset of nephropathy. Biopsies only provide information on the sampled region of the kidney, and do not capture the inherent heterogeneity of BKV infection within the kidney allograft and disease progression. The assay described here may find use as a noninvasive alternative to conventional biopsy to screen for BKV related kidney injury.

In addition, patients with bacterial UTIs show higher neutrophil contributions, suggesting immune activation and recruitment of neutrophils to the urinary tract, as well as elevated amounts of bladder cfDNA, indicating tissue injury. A common question in infectious diseases is how to interpret positive urine cultures. Outside of specific indications such as pregnancy, urological procedures, and being within 3 months of transplant^42,43^, a positive urine culture is currently treated with antibiotics only in symptomatic individuals. By quantifying the release of cfDNA from different cell types and tissues, the assay reported here can provide clinicians with additional information to guide treatment decisions.

A quantitative measurement of the tissues-of-origin of cfDNA constitutes a generalizable noninvasive approach to identify injury to vascularized tissues and participation of the immune system and can find wide application in diagnostic medicine. Several studies have reported technologies to trace the tissue origin of cfDNA in wide range of settings, targeting different epigenetic marks comprised within cfDNA, including the footprints of nucleosomes and transcription factors, cytosine methylation and hydroxymethylation ^7–9,44,45^. Here, we have applied this technology for the first time to the monitoring of host tissue damage due to infection. Host-based molecular signatures have previously been considered as classifiers of infectious complications. In respiratory infection, host transcriptional profiling of the peripheral blood has been shown to provide a means to characterize the host response to viral and bacterial infection, and to discriminate between infectious and noninfected states^46^. Host-response molecular signatures thereby offer a diagnostic approach orthogonal to approaches that focus on viral and bacterial pathogens. Recently, Langelier et al. described a diagnostic approach that combines transcriptional profiling and metagenomic sequencing of tracheal aspirate to classify lower respiratory tract infection^47^. Here, we show that a cfDNA assay can be used to screen for infectious agents and to quantify the degree of host damage. The strength of this assay lies in its simplicity of implementation, its noninvasiveness, and its ability to directly interrogate host damage, the metric that is most relevant to classify infectious complications in the framework proposed by Casadevall and Pirofski^5^.

In summary, we propose that WGBS of cfDNA can be used as a metagenomic sequencing assay to provide in depth understanding of both the metagenome as well as the host response to infection. This assay is generalizable to multiple diseased states and has the potential to distinguish colonization from infectious disease in a clinical setting.

## MATERIALS AND METHODS

### Study cohort

51 urine samples were collected from 36 kidney transplant recipients treated at NewYork-Presbyterian Hospital-Weill Cornell Medical Center. The study was approved by the Weill Cornell Medicine Institutional Review Board (protocols 9402002786 and 710009490 and 1207012730). All patients provided written informed consent.

Twenty-two urine samples from 21 patients were collected at the time of kidney allograft biopsy; 9 samples were collected from patients with BK viremia were found to have BK polyomavirus nephropathy with positive immunohistochemical staining for SV40 large T antigen (BKVN), 7 samples from patients with BK viremia were found not to have BK polyomavirus nephropathy (BKV+/N-) and 6 patients did not have BK viremia and had no significant pathology in the biopsies (Normal).

Plasma BKV copies per ml of blood were measured using the quantitative assay developed by Quest Diagnostics® as part of clinical testing.

### Renal Allograft Biopsy Evaluation

Percutaneous core needle biopsy specimens of kidney allografts were fixed in 10% buffered formalin and embedded in paraffin. Tissue sections were stained with hematoxylin eosin, periodic acid Schiff, periodic acid silver methenamine, and Masson trichrome for light microscopic evaluation. Routine immunofluorescence staining of fresh frozen biopsy tissue and immunoperoxidase staining of paraffin-embedded biopsy tissues were performed using standard techniques to detect the presence of positive staining for C4d (Fisher Scientific – C4d polyclonal NC9575575; Quidel – C4d monoclonal Cat. No. A213) and SV40 large T antigen of the BK virus (Affinity purified and agarose conjugated IgG2A mouse monoclonal antibody recognizing the 94kDa SV40 large T antigen; PAb416, Cat. No. DPO2, Calbiochem, USA) respectively.

Twenty-four urine samples were collected from patients who underwent same day urine culture^2^. Briefly, urine samples were plated on tryptic soy agar with sheep blood and incubated at 35C. Samples were classified as UTI positive (UTI, n=12) if an organism was detected at least at the genus level with 10,000 colony forming units (cfu) per ml. Samples were classified as UTI negative (no-UTI, n=12) if no organism was isolated (n=11), or if no organism was identified at the genus level and the colony counts was under 10,000 cfu/ml (n=1). Early time point (n=5) and Normal (n=6) samples were selected from patients who did not present symptoms for BK polyomavirus or UTI and who provided a urine sample within three days, or following 100 days after transplantation, respectively. Normal samples were collected from patients who underwent routine biopsy as part of standard clinical practice and did not show histological evidence of inflammation or nephropathy.

### Urine collection, supernatant isolation and cfDNA extraction

Urine was collected using a conventional clean-catch method (n=46) or through foley catheter (n=5). Approximately 50 ml of urine was centrifuged at 3,000g for 30 minutes on the day of collection and supernatants were stored at −80C. cfDNA was extracted from 1 or 4 ml of urine according to manufacturer recommendations (Qiagen Circulating Nucleic Acid Kit, Qiagen, Valencia, CA) and quantified using a Qubit fluorometer 3.0 (high sensitivity double-stranded DNA kit, Thermofisher, Waltham, MA). The concentration of cfDNA in the sample was calculated by multiplying the measured concentration of DNA in eluant by the elution volume and dividing by the volume of the urine sample.

### Bisulfite treatment, library preparation and Illumina sequencing

Between 5 and 20 µl of eluted cfDNA was bisulfite-treated using the Zymo EZ DNA methylation kit according to manufacturer’s recommendations (Zymo EZ DNA methylation kit, Irvine, CA). Samples were eluted in approximately 30 µl. Samples were prepared for sequencing using a single-stranded library preparation^16,17^. Libraries were characterized using DNA fragment analysis (Advanced Analytical Fragment Analyzer) and sequenced (Illumina NextSeq550, 2×75bp).

### Alignment to the human genome and methylation extraction

Low-quality bases and adapter sequences were trimmed (Trimmomatic v-036^48^). Reads were aligned to a C-to-T converted hg19 reference using bwa-meth v0.2.0^28^. PCR duplicates and low quality reads were removed using Samtools v1.16^49^. Methylation densities were measured using MethylDackel v.0.3.0-3-g094d926^50^.

### Quantification of urine-derived microbial cfDNA

The burden of microbial cfDNA after conventional sequencing was determined as previously described^3^. The burden of microbial cfDNA after WGBS was determined using a similar approach. Briefly, low-quality bases and adapter-specific sequences were trimmed (Trimmomatic^48^, v036) and short reads were merged (FLASH^51^ v1.2.11). Reads were aligned to a C-to-T converted hg19 reference using bwa-meth^28^ (v0.2.0) and to the standard hg19 genome using bwa^49^ (v.0.7.13-r1126) to remove converted and unconverted host reads. Reads were then BLASTed^52^ to a list of C-to-T converted microbial reference genomes, and a relative abundance of each organism was determined using GRAMMy^40^. The fraction of reads of microbial origin was defined as the ratio of reads mapping to a microbial reference to the total number of paired-end reads for that sample.

### Bisulfite conversion efficiency

Although CpG dinucleotides are often methylated in the human genome, C[A/T/C] molecules are rarely methylated. We estimated bisulfite conversion efficiency by calculating the reported rate of C[A/T/C] methylation using MethPipe v.3.4.3^53,54^.

### Donor fraction measurement

Donor-specific cfDNA was measured in sex-mismatched donor-recipient pairs as previously described^2^ by first adjusting sequence mappability with HMMcopy^55^.

### Cell and tissue methylation reference preparation

References made available by public consortia were downloaded (Supplemental table 1, refs). Genomic coordinates of references aligned to hg38 assembly of the human genome were converted to the hg19 assembly using CrossMap^56^. Single base pair CpG methylation values (forward strand) were extracted, and all references were merged by genomic coordinates using BEDTools^57^. Methylation profiles were grouped by tissue-type, and differentially methylated regions (DMRs) were found using approaches implemented in Metilene (difference >= 20%, q-value < 0.05)^23^ in a one-versus-one approach. Overlapping regions were merged, and tissue methylation profiles were averaged over those regions. Regions were constrained to having a minimum of 10 CpGs per regions, with at least 1 CpG per 100 base pairs.

### Tissues-of-origin deconvolution

Tissues and cell-types of origin were determined by quadratic programming^58,59^ according to the following equation:

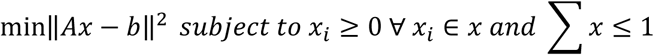

Where A is an (*m*+1) × *n* matrix with m cell and tissue methylation references spanning *n* DMRs. The additional column contains an error parameter to consider potential missing references, non-tissue specific methylation, and other sources of error. x_i_ represents the contribution of each tissue to the cfDNA mixture. b is a 1×n vector of the observed methylation. Only autosomal chromosomes were considered for tissue of origin measurements. The absolute concentration of tissue specific cfDNA in the samples was calculated by multiplying the tissue proportion by the absolute cfDNA concentration of the sample and by the fraction of sequenced reads of human origin.

### Dimensional reduction

Dimensional reduction was performed on DMRs selected in the reference that were present in all samples. Clustering was performed using PCA (R, prcomp package) and UMAP (R, umap package). Default UMAP parameters were used (15 neighbors, 2 components, Euclidean metric and a minimum distance of 0.1). Sample methylation profiles were projected onto the two-dimensional feature spaces of PCA and UMAP using the predict function in R.

### Microbe detection through sequencing

For each culture-detected microbe (*E. coli, E. faecalis, E. faecium, P. aeruginosa, K. oxytoca and coagulase-negative Staphylococcus*), a specific threshold corresponding to the smallest relative genomic abundance with positive culture was used to classify all tested samples. Samples were classified as being positive (RGE [tested microbe] ≥ minimum threshold [tested microbe]) or negative for each culture-detected microbe (RGE [tested microbe] < minimum threshold [tested microbe]), and a confusion matrix was used to compare predicted values through sequencing with reference values determined by urine culture. Test accuracy statistics were calculated using R (caret package). Samples were classified as virus and/or bacteria positive if the relative genomic of a microbe was greater than set thresholds (10^3^ for BK virus, 0.09 for bacteria) and if the microbe is not a known sequencing contaminant^60^ (common sequencing contaminants that are also known uropathogens were classified as positive).

## Statistical analysis

All statistical methods were performed in R (v.3.5). All groups were compared using a two-sided Wilcoxon test (R stats package). Boxes in the boxplots indicate the 25th and 75th percentiles, the band in the box indicates the median, lower whiskers extend from the hinge to the smallest value at most 1.5× IQR of the hinge, and higher whiskers extend from the hinge to the highest value at most 1.5× IQR of the hinge. Violin plots are bound by density estimates of the data distribution.

## Data availability

The sequencing data generated for this study will be made available in the database of Genotypes and Phenotypes (dbGaP).

## Code availability

All custom scripts are available at https://github.com/alexpcheng/bisulfite_cfDNA

## Materials and Correspondence

All requests should be submitted to I.D.V. and D.D.

## Supplementary Materials

**Figure S1**. Principal component analysis of reference methylomes.

**Figure S2**. Scatterplot of corresponding donor fractions in sex-mismatched samples that underwent both WGBS and standard sequencing.

**Figure S3**. Mean fragment length of bisulfite-treated cfDNA for BKVN, BKV+/N-, Normal, no-UTI, UTI and Early groups.

**Table S1**. Reference methylation dataset metadata

**Table S2**. Clinical information of urine samples

## AUTHOR CONTRIBUTIONS

A.P.C., P.B., M.P.C., J.R.L, D.D., M.S., and I.D.V. contributed to the study design. A.P.C. performed the experiments. A.P.C., P.B., J.R.L., D.D., and I.D.V. analyzed the data. A.P.C., M.P.C., D.D., J.R.L., and I.D.V. wrote the manuscript. All authors provided comments and edits.

## ACKNOWLEDGMENTS

We thank Peter Schweitzer and colleagues at the Cornell Biotechnology Resource Center (BRC) for help with sequencing assays. This work was supported by US National Institute of Health (NIH) grant 1DP2AI138242 to IDV, US NIH, grant 1R21AI133331 to IDV and JRL, a National Sciences and Engineering Research Council of Canada (401236174) fellowship to APC, and a National Science Foundation Graduate Research Fellowship Program (NSF-GRFP) grant DGE-1144153 to PB.

## Supplemental Materials for

**Supplemental figure 1.**
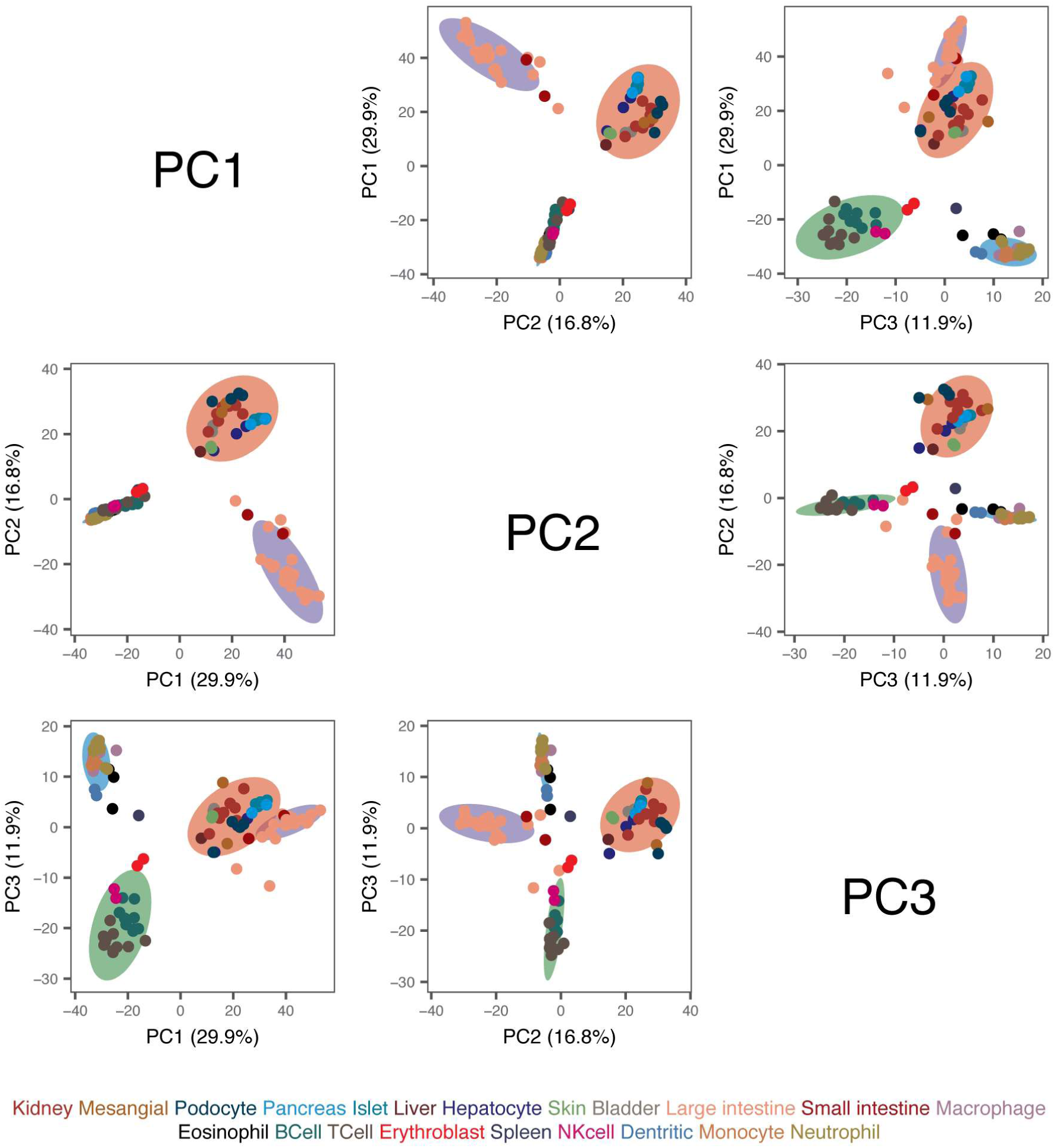
Principal component analysis of reference whole-genome methylation profiles from human tissues.

**Supplementary figure 2.**
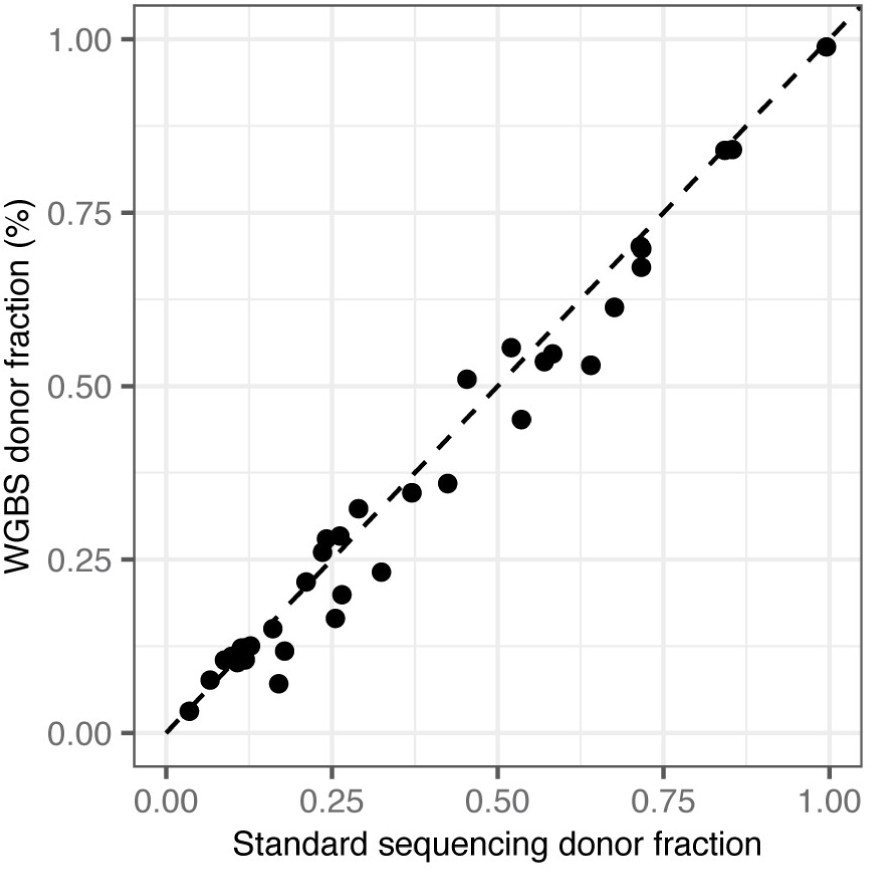
Comparison of proportion of donor-derived urinary cfDNA measured by conventional and whole-genome bisulfite sequencing (WGBS, n=36 matched samples). Dashed line is a one-to-one slope. Spearman’s rho = 0.97, p-value < 2.2×10^-16^.

**Supplemental figure 3.**
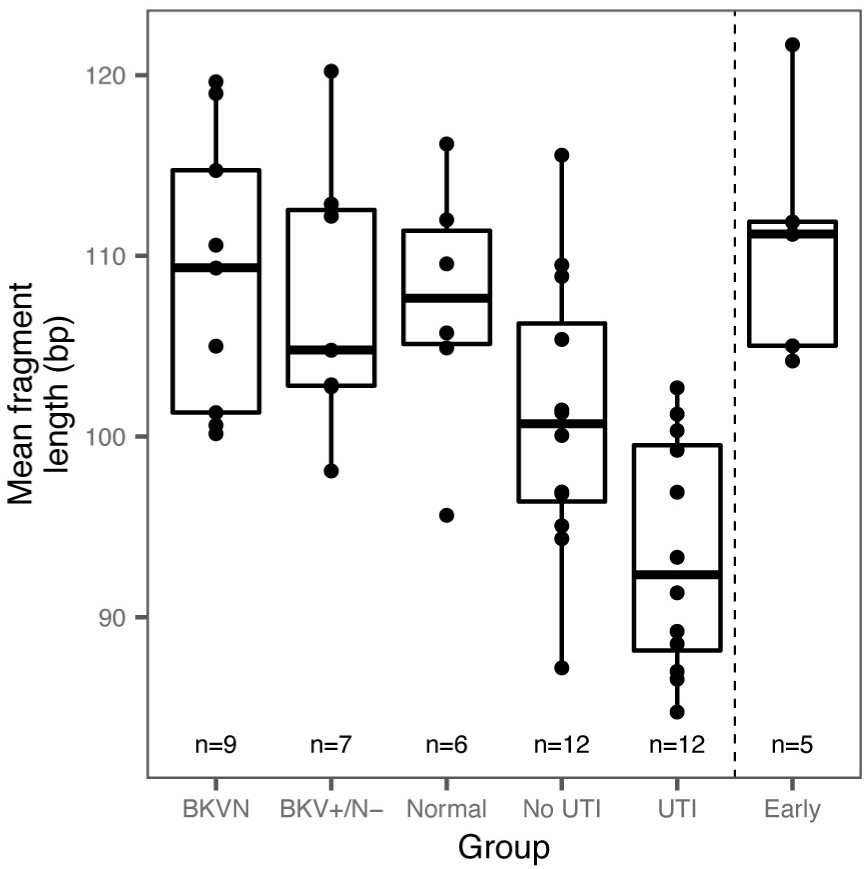
Mean fragment length of bisulfite-treated cfDNA for BKVN, BKV+/N-, Normal, no-UTI, UTI and Early groups (n=51).

